# Assessing Protein Homology Models with Docking Reproducibility

**DOI:** 10.1101/2022.02.17.480964

**Authors:** Alexander P. Plonski, Scott M. Reed

## Abstract

Results of the most recent Critical Assessment of Protein Structure (CASP) competition demonstrate that protein backbones can be predicted with very high accuracy. In particular, the artificial intelligence methods of AlphaFold 2 from DeepMind were able to produce structures that were similar enough to experimental structures that many described the problem of protein prediction solved. However, for such structures to be used for drug docking studies requires precision in the placement of side chain atoms as well. Here we built a library of 1334 small molecules and examined how reproducibly they bound to the same site on a protein using QuickVina-W, a branch of the program Autodock that is optimized for blind searches. We discovered that the higher the backbone quality of the homology model the greater the similarity between the small molecule docking to the experimental and modeled structures. Furthermore, we found that specific subsets of this library were particularly useful for identifying small differences between the best of the best modeled structures. Specifically, when the number of rotatable bonds in the small molecule increased, differences in binding sites became more apparent.

## Introduction

Protein structure prediction has been an important scientific challenge in computational biology for many decades.^1^ Scientists have utilized molecular dynamics in order gain insight into protein folding especially for small proteins^1–3^ however molecular dynamics simulations are computationally expensive for medium to large proteins.^4, 5^ Homology modeling has been a relatively fast tool to get a predicted proteins structure based upon experimental structures that exist in the Protein Data Bank (rcsb.org), no matter the size of a protein.^6, 7^ Homology modeling has been limited by the templates of experimental structures available, and the quality of the model is affected by the evolutionary distance between the target and template proteins.^6^ AlphaFold is a deep-learning approach to homology modeling based on a trained neural network. The most recent version of this software, AlphaFold 2, demonstrated very high accuracy in predicting protein structures in the 14^th^ Critical Assessment of Protein Structure (CASP14).^8–10^ AlphaFold 2 was then used to produce predicted structures for an estimated 98% of the human proteome with similar coverage for many other organisms.^11^

Molecular docking has been a popular tool in computer-aided drug discovery. It is a computationally inexpensive method for understanding drug-protein interactions, including a drug’s binding conformation and affinity.^12^ While molecular docking can provide useful insight into molecular interactions, it is unknown how accurate docking studies are with homology models on the molecular level. Results from using homology models for docking are mixed.^13^ Previous work has been done in benchmarking the use of homology models with protein-protein docking which has shown the general utility of these models based on their sequence similarities and C_α_ Root Mean Square Deviations (RMSD) from experimental structures.^14^ This approach has not provided much information on the molecular level details of a protein model.

While AlphaFold 2 does not elucidate the physical manner in which a protein folds,^15^ it has been shown to predict protein backbones with high accuracy when compared to experimental structures, however, less attention has been paid to side-chain structure in AlphaFold 2 models. A recent paper reported that for the three CASP14 proteins that had bound ligands of *S*-adenosylmethionine or adenosine-5’-diphosphate in their experimental structures, the homology model had the same location of binding but the binding pose was different than the experimental structure.^16^ These molecular details are crucial to protein function and specifically important in studies of drug-protein interactions, where the misidentification of a side chain in a binding site could potentially affect the accuracy in modeling binding of drugs.

Both the methods for preparing AlphaFold structures and the method for evaluating their success in CASP are focused on the three-dimensional positions of backbone atoms. CASP evaluation is based on the similarity of backbone atoms to experimental structures, ignoring side chains. The Global Distance Test Total Score (GDT_TS) is used to evaluate CASP entries which is the average of the percentage of residue backbone atoms within 1, 2, 4, and 8 Å of the experimental structures. AlphaFold first made its appearance in CASP13, where it had the highest accuracy of the competitors, with a median GDT_TS of 62.12.^17^ Since then, DeepMind has updated their neural network architectures and training procedures and achieved a median GDT_TS of 89.42 in the CASP14 competition with AlphaFold 2.^9, 10^ AlphaFold predictions add side chains to the structures after artificial intelligence is used to construct the three-dimensional arrangement of the peptide backbone.^9^ The optimization of the side chains is performed using the Amber forcefield as a final step in the workflow and does not involve the use of artificial intelligence.^11^

Each of the residues in the publicly available AlphaFold 2 proteome structures are assessed with a predicted Local Distance Difference Test (pLDDT) score between 0 and 100 for each residue that is a prediction of the Local Distance Difference Test (LDDT) for C_α_ atoms. The LDDT is a superposition-free scoring function used to evaluate local distance differences and calculated similarly to the GDT_TS as an average of fraction of atoms within a threshold of 0.5, 1, 2 and 4 Å of their comparitor.^18^ Being able to connect pLDDT scores to potential for accuracy in docking studies will facilitate the accurate use of homology models and avoid errors resulting from docking to poor quality models.

Here we examine the accuracy of the side chains in the AlphaFold 2 structures and how their side chain structures impact the utility of the models for small molecule docking studies. We specifically examined the protein set that was evaluated in CASP14 which allowed for a comparison of AlphaFold structures to experimentally verified structures that were hidden from contestants during CASP14. Recent work has been done on assessing the binding properties of CASP14 models through a computational solvent mapping analog with a small number (16) of small molecule probes.^16, 19^

AlphaFold models have already demonstrated utility in molecular replacement methods for solving structures from x-ray structures.^20^ Docking studies, commonly used in drug design have a more stringent need for accurate models to provide meaningful data. Here we find that subtle difference between even the highest scoring AlphaFold 2 structures and their experimental counterparts can be revealed through docking. And large and flexible molecules are particularly well-suited for identifying these differences.

## Methods

### Selection and Preparation of Proteins and Molecules

AlphaFold structures and their corresponding experimental structures were obtained from the CASP-14 prediction database (https://predictioncenter.org/). Targets included both the AlphaFold prediction and experimental structure in the CASP-14 prediction database. Experimental structures from CASP14 were resolved via X-ray crystallography and solution NMR (only T1027 and T1029). The AlphaFold models with the highest GDT_TS were selected for docking. Targets with multiple unresolved atoms in the experimental structures were removed. Terminal regions in which residues were unresolved in the experimental structures were removed from the predicted structures, so that all residues were matched in sequence to their target. A total of 29 sets of experimental and modeled proteins were selected for the study.

Molecular probes for docking were obtained from human, yeast, and *e. coli* metabolome databases.^21–23^ They were filtered for a molecular weight range of 300 – 800 g/mol. Molecules containing selenium, iron, and molybdenum were removed. A total of 1334 molecules were selected for the study with 971, 38, and 325 coming from the human, yeast, and *e. coli* metabolome databases, respectively. 3D structures were generated using rdkit’s AllChem module,^24^ and charges at physiological pH were calculated using JChem’s cxcalc.^25^

### Molecular Docking

The MGLTools Package in AutoDock Tools was used to prepare target structures and molecules for docking using the prepare ligand and receptor scripts.^26^ QuickVina-W 1.1 was used for rigid molecular docking with a grid box size extending 10 Å beyond the protein in each dimension and an exhaustiveness search of 10.^27^ Docking studies were run on all experimental and AlphaFold structures on an HPE ProLiant ML110 G10 with 16 Intel Xeon cores. The highest scoring docked molecule for each structure was selected for analysis.

### Side Chain RMSD

The Multiscale Modeling Tools for Structural Biology (MMTSB) was used to calculate side chain RMSD between AlphaFold and experimental structures.^28^ A least squares fit was performed to align C_α_ atoms and C_β_ atoms carbons between structures within MMTSB prior to calculating side chain RMSD.

### Distance Measurements and Normalization

Distance measurements for cross (experimental-AlphaFold) and self-comparisons (experimental-experimental or AlphaFold-AlphaFold) were computed based upon the closest distance between the center of mass of a molecule and the closest alpha carbon on a residue of the first target structure. Similarly, distances between molecule docking results for the second structures were computed based upon the center of mass of a ligand and the C_α_ atom selected in the molecule-target structure distance computation. Distance measurements were normalized for each protein by taking a distance measurement and dividing it by the greatest distance between two atoms on that protein. The Python package Biopandas was utilized for loading protein and molecule structures into the dataframes format and calculating distance changes.^29^

### Measuring the Close-Binding Fraction of Molecules

The close-binding fraction of molecules for each protein was calculated by taking the count of distance measurements below 2 Å (using the method above) and dividing by the total number of ligands (1334).

### Molecule Intersects

Molecule intersects were calculated for molecules in the close-binding fraction of the top four GDT_TS-scoring proteins (T1056, T1046s1, T1035, and T1046s2). Molecular properties (tanimoto coefficient, molecular weight, logP, and rotatable bonds) of molecule intersects were calculated using rdkit.^24^ Distance measurements for molecules with the number of rotatable bonds equating to 5 and 9 were filtered and the mean of median distance changes across three trials for cross and self-comparisons were plotted along with their standard deviations.

## Results

### Performance of AlphaFold Structures in Docking Studies

First, we examined the accuracy of side chain placement in the CASP14 AlphaFold models, ignoring backbone atoms by using the Multiscale Modeling Tools for Structural Biology (MMTSB) package.^28^ We measured the RMSD values exclusively for the side chain atoms of the AlphaFold models compared to the corresponding experimental structure which allowed us to evaluate how closely the side chain atom positions match the experimental structures (Figure 1). The variability amongst the structures is high (3.75 ± 2.64 Å) and structures with higher GDT_TS backbone scores have lower RMSD values for the side chains, suggesting that more accurate backbones constrain the options available for the side chain placement and optimization by Amber such that they end up closer to their corresponding experimental structures. Importantly, structures T1027 and T1029 which have the highest side chain RMSDs (15.14 Å for T1027 and 9.11 Å for T1029) are NMR based experimental structures. Because the training set for AlphaFold models was based primarily on crystal structures, it is not surprising that the side chain placements differ.

**Figure 1.**
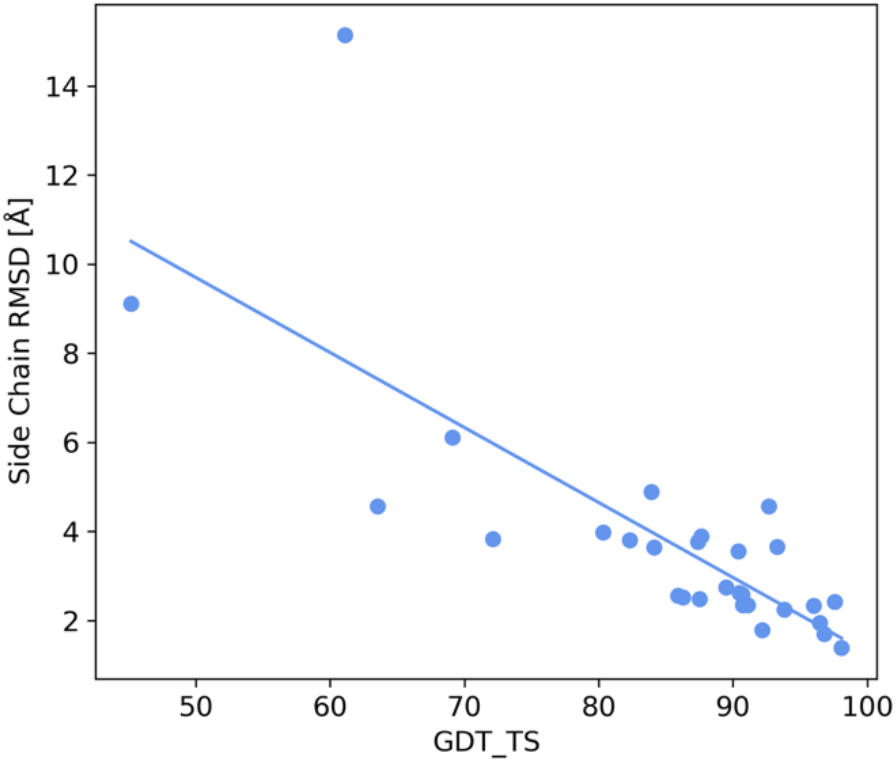
Scatter plot of AlphaFold model score (GDT_TS) and sidechain RMSD. The Pearson correlation coefficient is −0.77.

Only a few CASP14 proteins have known ligands but many molecules will bind promiscuously to proteins, so we created a library of molecules in hopes that some would find a suitable location to bind with high reproducibility. A similar approach of docking very small molecular probes to CASP14 structures demonstrated consistency in probe biding site but weak correlation to GDT_TS scores.^16^ Here the goal was not to measure binding affinity or to identify a known binding site but rather to find molecules that docked reproducibly at the same location for a given protein. The metric used was reproducibility using docking with non-native ligands to observe how homology models differ from experimental structures as receptors. To accomplish this, we created a set of 1334 molecules to use for docking. The molecules were selected from metabolite databases from multiple organisms but have no specific connection to the CASP14 proteins or the organisms from which the CASP14 proteins were sourced. But they contain a structurally diverse set of biologically relevant molecules.^21–23^ Each molecule was docked on both the experimental and AlphaFold protein structures for each of the target proteins using QVina-W with no constraints on the number of rotatable bonds. QVina-W is optimized for such blind searches without a predetermined cavity. The grid size was extended 10 Å greater than the protein dimensions ensuring a truly blind search, without biasing toward any pre-selected location.^27^ Without expected binding cavities, a cavity pre-screen was not necessary as is often done in large scale blind docking studies.^30^ A schematic of the process can be viewed in Figure 2.

**Figure 2.**
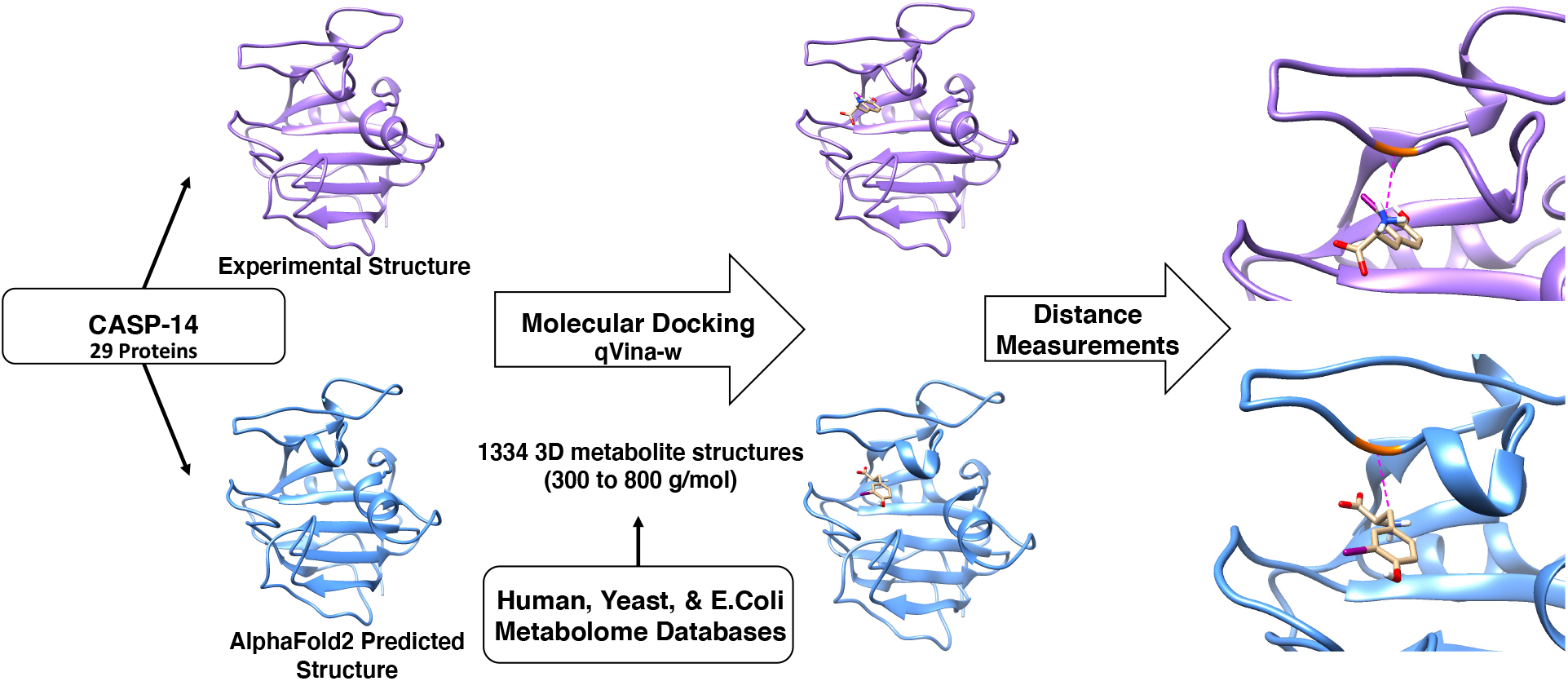
Selected molecules from metabolome databases are docked onto CASP-14 experimental and predicted protein structures, in which the distance between the same two contacts (molecule’s center of mass and protein’s closest C_α_ atom in the experimental) is calculated between experimental and predicted structures for each docking result.

Three trials of the docking study were repeated under identical conditions and for each docking trial the distance of the closest contact between the bound molecule’s center and the protein’s closest C_α_ atom was measured. Changes in this measurement to the same residue were then used to compare two different docking trails and multiple comparisons were made to understand the reproducibility of the docking poses. First, the distance changes between repeated trials for all combinations of two docking studies conducted on experimental structures (Exp-Exp, Figure 3, green) were examined followed by all combinations of two docking studies conducted on AlphaFold models (Af-Af, Figure 3, red). The distance distributions for these self-comparisons were plotted as box and whisker plots for each protein. In all proteins, regardless of which two trials are compared there is consistency in the interquartile range (IQR) of distances for each protein showing the reproducibility of this approach. In general, the IQR was larger for proteins with lower GDT_TS values. A few proteins such as T1030 and T1050 show particularly high distributions of distance changes in the self-comparisons. This is due to the size of the proteins and that these proteins bound molecules inconsistently, allowing for variability between trials.

**Figure 3.**
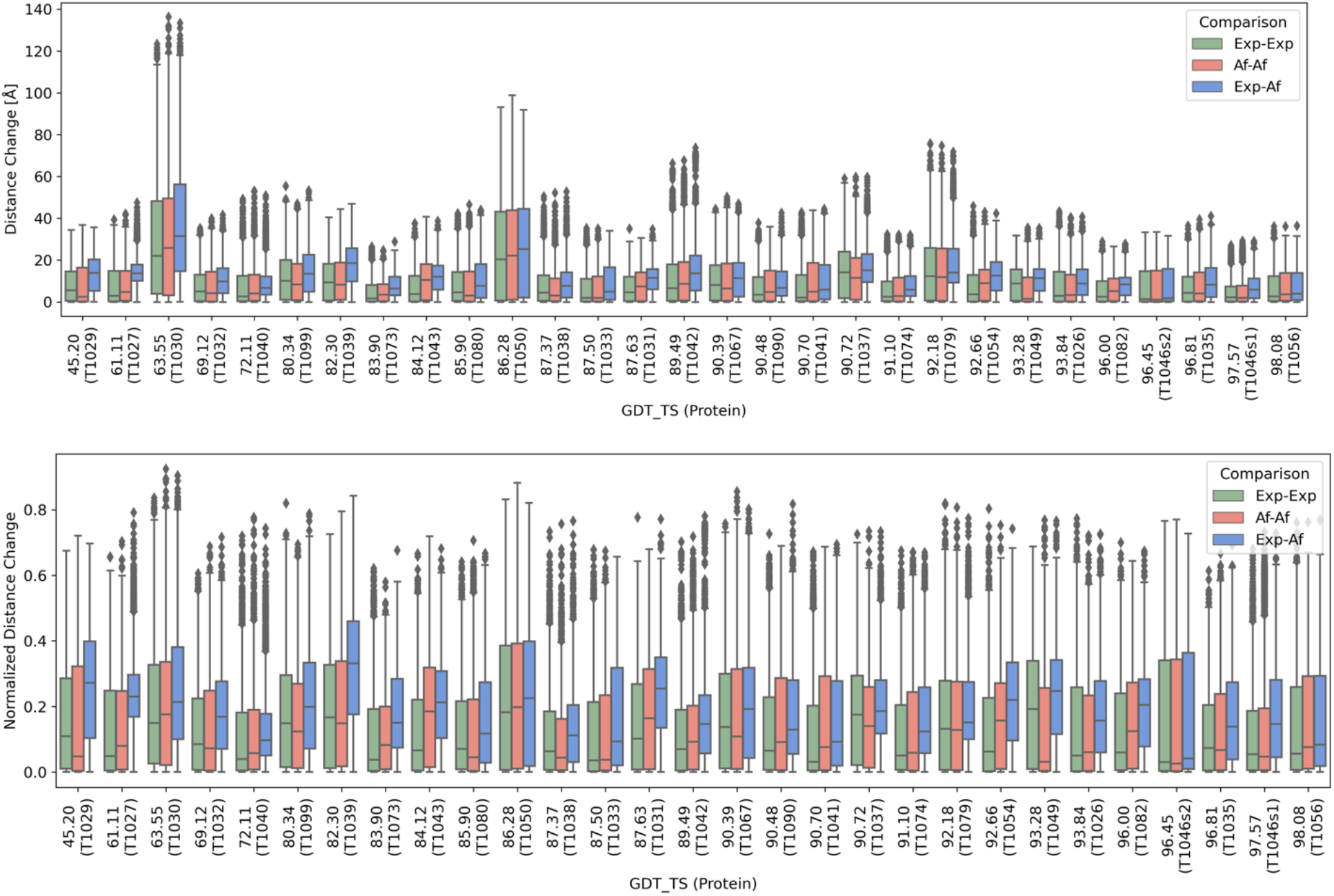
Comparison of distance changes for repeated docking to the same structure (Red = AlphaFold, Green = experimental) and distance changes between docking to AlphaFold structure compared to docking to experimental (Blue). Actual distance (top) and normalized distance (bottom).

CASP14 proteins ranged in size from 8 to 100 kDa and one clear trend in the data is that larger proteins have a greater range of possible distance changes, as expected. To account for the range in protein size, the data was normalized for each protein to the range of possible distances across the surface of that protein, such that a value of 1 corresponds to the greatest distance between two atoms on the protein and 0 corresponds to no change in distance between the two protein structures. These normalized distance distributions were plotted both as boxplots and as violin plots with the kernel density displayed on the vertical axis for each protein (Figure S1). Again, heterogeneity in the docking results is observed for different proteins without a strong trend in the positions of the IQR as a function of GDT score.

Next, we analyzed the difference between the highest ranked pose of the docked molecules between the AlphaFold and experimental structures (Exp-Af, Figure 3, blue). We compared this to the distance between these same two contacts in the AlphaFold model to obtain a distance change for all 1334 molecules for each of the proteins. The distance distributions for this cross comparison are also displayed as box and whisker plots for each protein in Figure 3. Overall, there is a similarity between the boxplots for self-comparisons and for the boxplots that compare docking to experimental structures and AlphaFold structures (cross-comparisons). The size of the IQR is roughly similar between the cross-comparisons and the self-comparisons and the number of outliers and on which proteins they occur is similar. There is a small but noticeable increase in the IQR for the cross comparisons especially for lower GDT value proteins. This suggests that the poorer the model quality, the less consistency there is between docking to the model and docking to the experimental structure. However, there is not a general broadening of the IQR when looking at cross comparisons compared to self-comparisons, suggesting the homology models are reasonable surrogates for the experimental structures.

In both the self- and cross-comparisons, the median values for distance changes are consistently below the mid-point in the range of distance measurements possible for a protein whereas if the docking was not identifying binding sites consistently, the molecules would be placed randomly, and therefore the median values would be slightly larger than the radius for a spherical, globular protein.^31^ In general, the first three quartiles of distance changes are below what would be expected for random placements. This same data for Exp-Af is also plotted as violin plots with the kernel density displayed on the vertical axis for each protein to show the distribution of distances more clearly. In most cases a cluster of very short distances is noticeable, and this is noticeable when to the data is averaged over all trials (Figure 4) or when examining individual trials (Figure S2).

**Figure 4.**
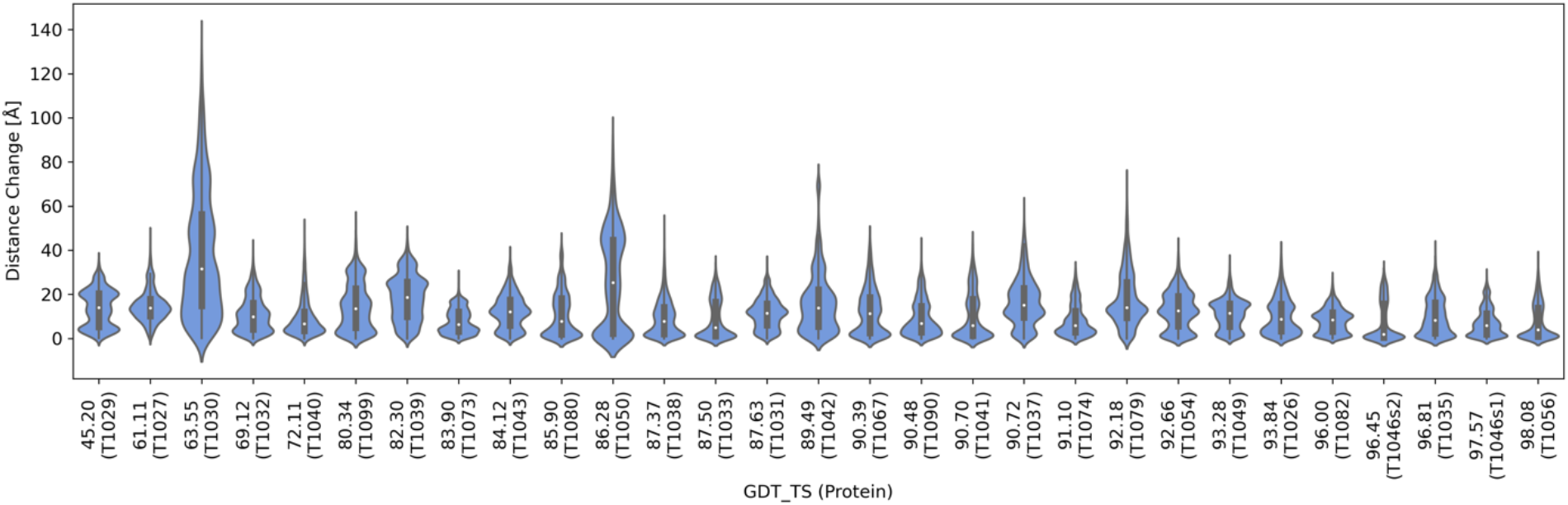
Distribution of non-normalized distance changes for docked molecules between experimental and AlphaFold models.

The distribution of distances and kernel density for cross comparisons (Figure 4) provided a unique fingerprint for each protein and this distribution of distances was very reproducible for proteins with a high GDT_TS (Figure S3). Proteins with a higher GDT_TS show a lower cross comparison median when compared to low GDT_TS models. Similarly, as in the self-comparisons, proteins T1030 and T1050 show a higher distribution of distance changes than the proteins determined by x-ray.

The differences in distance values of most interest are those that could be considered to reveal the molecule is staying within the same binding site. Since these proteins are not expected to have a natural cavity for any of the molecules being docked, the very close distance cut-off (within 2 Å) between the experimental and AlphaFold structures was used to define two molecules as being in the same binding site. The higher GDT_TS proteins have a higher fraction of the 1334 molecules within this 2 Å cutoff. There is a modest trend in the fraction of molecules contained within this cutoff for self-comparisons; a 0.28 Pearson correlation coefficient for experimental structure and 0.17 for AlphaFold structures (Figure 5). There is a much stronger Pearson correlation coefficient of 0.57 for cross comparisons. This suggests that a very high GDT_TS is required for a protein to serve as a good model for docking studies. The modest correlation of GDT_TS to the distance changes for the self-comparisons of AlphaFold structures may reflect the general compactness of the structures which in turn could influence reproducibility of the docking. The fraction of molecules within 2 Å for cross-comparisons approaches that of the self-comparisons but does not meet it, showing that the models are not identical to their experimental counterparts as a receptor for docking.

**Figure 5.**
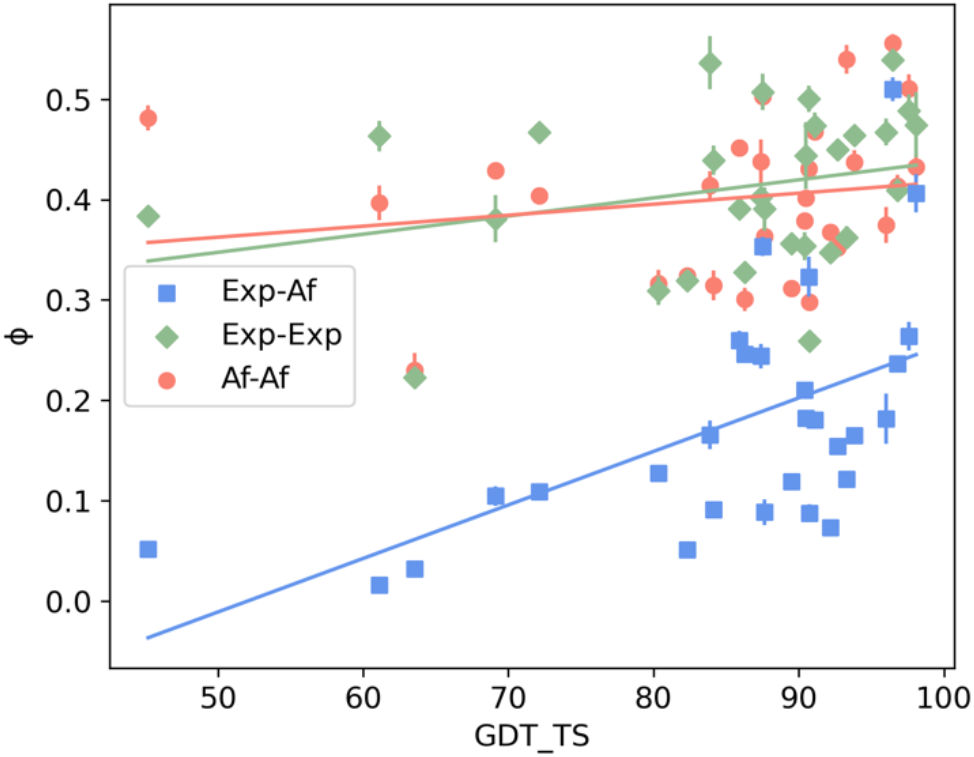
Scatter plot of AlphaFold model score (GDT_TS) vs the close binding fraction (phi) for cross- and self-comparisons. The Pearson correlation coefficients are: 0.57 for Exp-Af, 0.28 for Exp-Exp, and 0.17 for Af-Af.

We tested an alternate explanation for this GDT_TS trend by examining the correlation between protein size and the fraction of small molecules that were within the 2 Å cutoff (Figure S4). The correlation was much weaker than the correlation to GDT_TS and was weakest for the cross-comparisons (r = −0.10), suggesting that protein size is not an explanation for the observed trend between GDT_TS and docking change similarity. Molecular weight was in fact more strongly correlated to the fraction within the 2 Å for the self-comparisons (r = −0.49), suggesting that in those cases the larger proteins were more likely to possess multiple binding sites of similar character thereby reducing the number of molecules withing 2 Å.

The distribution of distances provided a unique fingerprint for each protein and the distribution of distances was very reproducible for proteins with a high GDT_TS. In all proteins studied, the highest normalized distance value for a molecule was 0.89 (for structure T1030) and the mean high value across all 29 proteins was 0.69. Given that a value of 1 would require a molecule to be in close contact with two atoms on the two most distal regions of a protein in the two structures, it is not surprising that the distributions approached but did not reach 1. Being in close contact with these outlier atoms would likely involve minimal contact with the protein, resulting in a low score and that pose not being ranked highly.

### A Unique Subset of Molecules are Highly Reproducible in Docking

Next, we examined what molecular characteristics were important to making molecules likely to appear in the same location on repeated trials for the same experimental structure (Exp-Exp), the same homology model (Af-Af), or the same location between experimental and homology models (Exp-Af) (Table 1). These are specifically the subset of the original 1334 molecules that bind within the close binding fraction (under a 2 Å distance change) in the four top scoring protein structures (T1056, T1046s1, T1035, and T1046s2). If docking always produced identical results, these intersects would include all 1334 molecules. Deviations from this number reveal the promiscuity of a given probe and the uniqueness of binding sites in those four proteins. The total number of molecules in this intersect is higher (~250) for the self-comparisons than for the cross comparison (~30). In both, the variability in trials is small expect for logp. The properties of these subgroups differ from the bulk with the molecular weights, logp values, and number of rotatable bonds all being lower than those values in the larger set of 1334 molecules. The Tanimoto coefficient which ranges from 0 to 1 and measures the similarity between two molecules based upon a molecular fingerprint identity was also examined.^32^ The similarity between molecules within each group, as expressed by the average point-wise calculation of a Tanimoto coefficient for each pair, is also very similar.

**Table 1.**
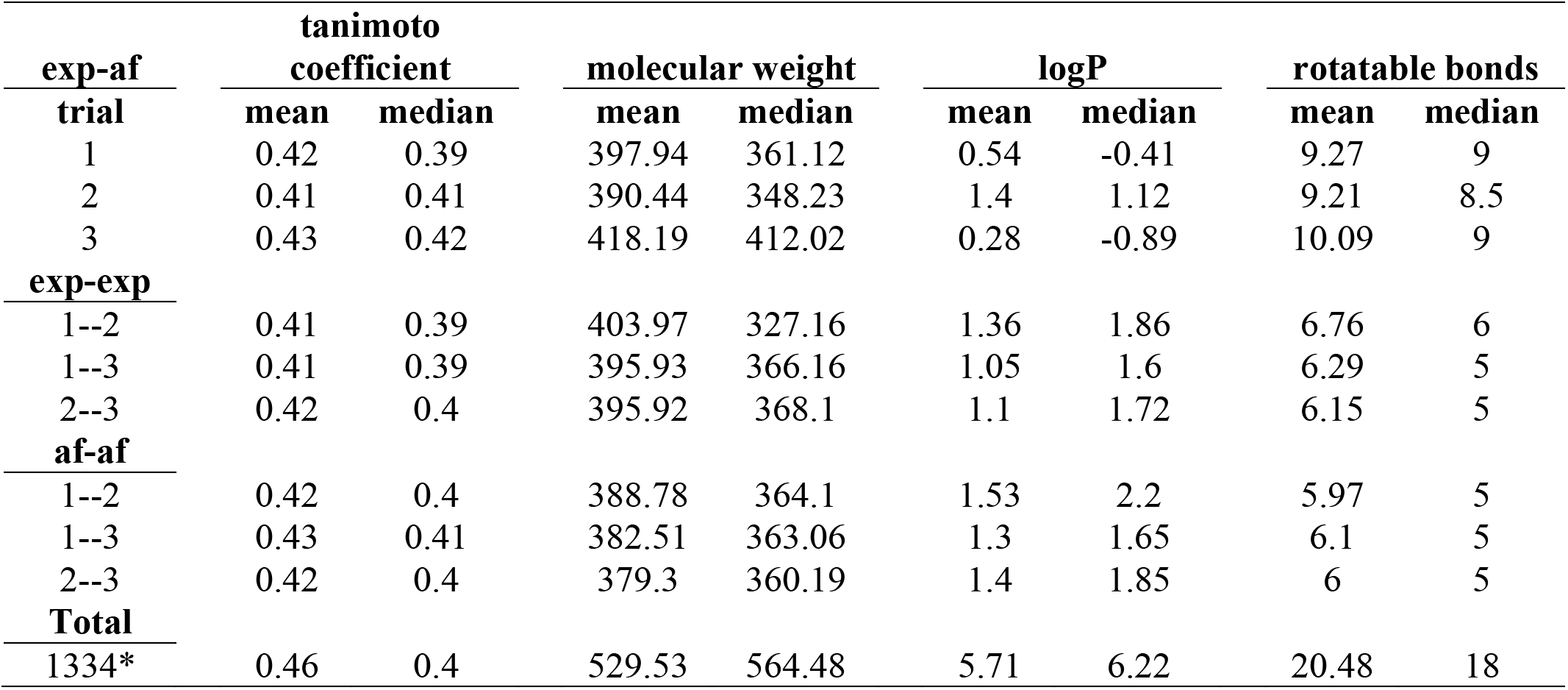
Mean and median values for properties of molecules present in all the top four scoring AlphaFold 2 proteins (T1056, T1046s1, T1035, and T1046s2) below a 2 Å distance change in docking trials 1, 2, and the full set of 1334 molecules. Trials 1 and 2 had 30 and 28 molecules respectively.

We next looked at properties of the molecules to see if they differed between self-comparisons (middle and bottom groupings, Table 1) and cross-comparisons (top grouping, Table 1). Most properties, including molecular weight, logp, and tanimoto coefficients are not very different between any of the subgroups. One property however did vary. Molecules that appeared in the intersect of cross-comparisons have a significantly higher number of rotatable bonds (9) than the those in the intersects of self-comparisons (5) (molecular structures in Figures S5 and S6).

We then re-examined the docking data to look at the fraction of molecules that bound within 2 Å between trials, limiting the analysis to the subset of molecules from the larger group of 1334 that had either 5 (n = 54) or 9 (n = 42) rotatable bonds (Figure 6). The difference between the self-comparisons and cross-comparisons were more clearly visible in both analyses. For probes with 5 rotatable bonds, the self-comparisons had IQRs near zero while the cross-comparisons had broader ranges of distance changes. For the molecules with 9 rotatable bonds, the self-comparisons had higher IQRs than for 5 rotatable bonds although they were almost always lower than the IQR for the cross-comparison. Furthermore, examining the root mean square displacement (RMSD) to compare differences in the placement of the molecules using LS-Align^33^ revealed a greater difference between the self- and cross-comparisons when the probes had 5 rotatable bonds as compared to those with 9. (Figure S7)

**Figure 6.**
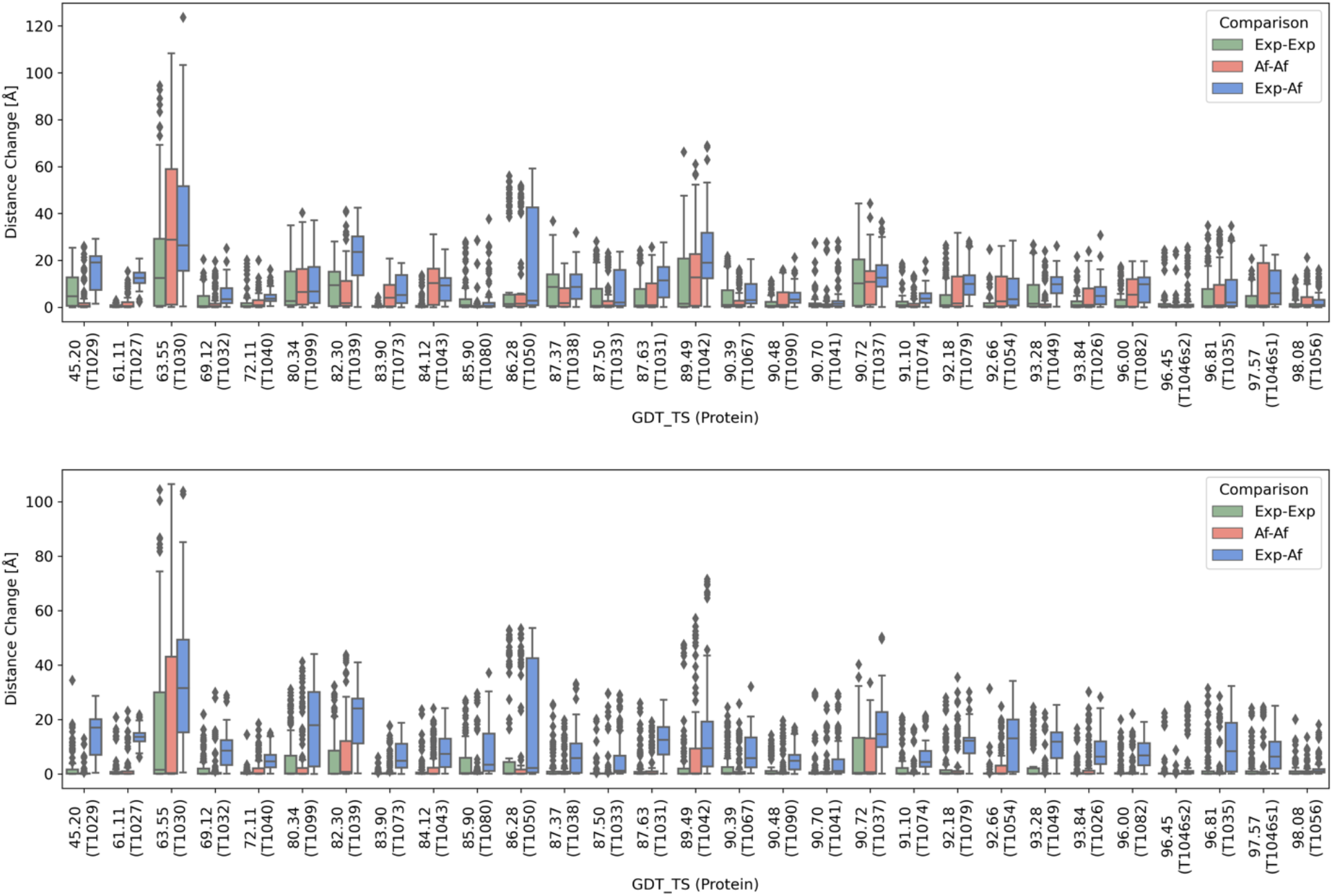
Comparison of distance changes for repeated docking to the same structure (Red = AlphaFold, Green = experimental) and distance changes for docking to AlphaFold compared to docking to experimental (Blue). Subset of docking molecules that have 9 rotatable bonds (n = 42) (top) or 5 rotatable bonds (n = 54) (Bottom).

Then looking at the median distance change for these subsets of molecules (Figure 7) the correlations with the GDT_TS score become much more apparent. The self-comparisons again show very little dependence on GDT_TS score. The slope for Exp-Exp was −0.073 and for Af-Af was −0.013 for molecules with 9 rotatable bonds. For molecules with 5 rotatable bonds, the slope for Exp-Exp was −0.00026 and Af-Af was 0.00113. In contrast, the cross-comparisons now reveal a strong trend with a slope of −0.243 for molecules with 9 rotatable bonds and −0.189 for molecules with 5 rotatable bonds and Pearson correlation coefficients of −0.98 and −0.96, respectively. Both subsets of molecules reveal trends that were masked in the larger set of 1334 molecules.

**Figure 7.**
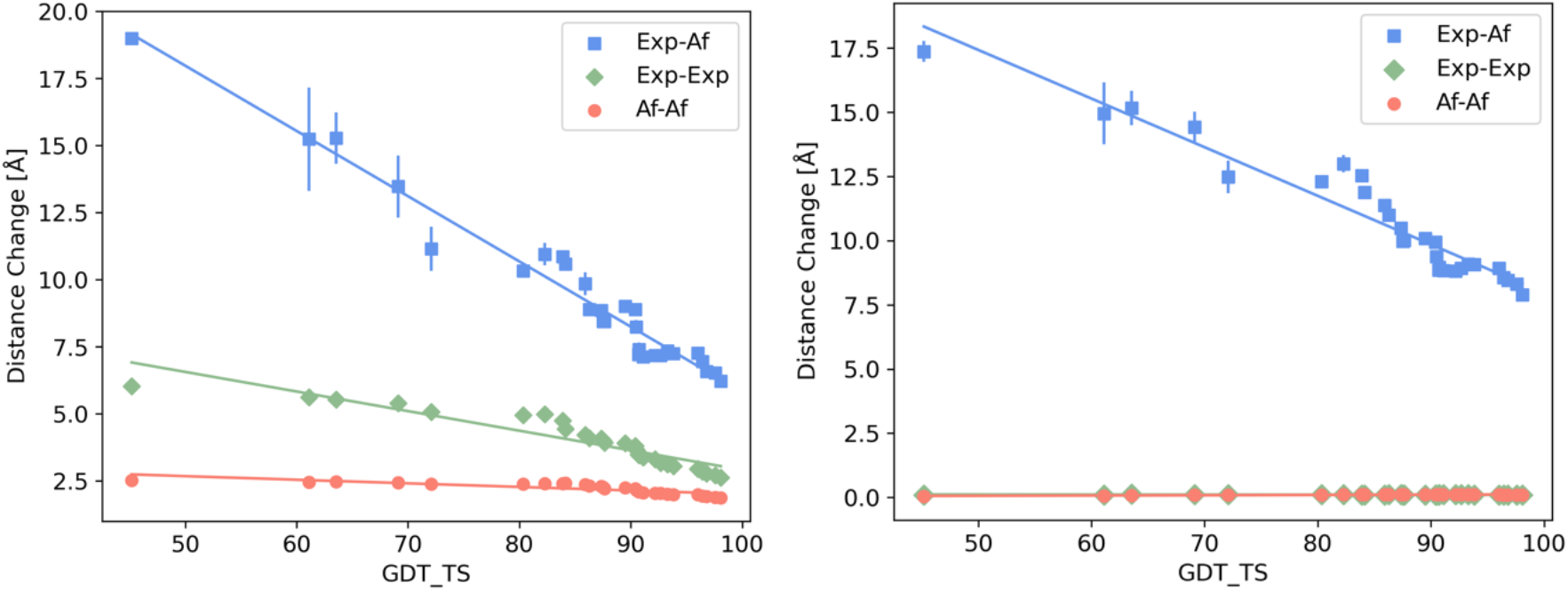
Comparison of median distance changes for repeated docking to the same structure (Red = AlphaFold, Green = experimental) and median distance change for docking to AlphaFold compared to docking to experimental (Blue) as a function of protein GDT_TS score. Subset of docking molecules that have 9 rotatable bonds (n = 42) (left) or 5 rotatable bonds (n = 54) (right). The Pearson correlation coefficients are as follows: Left: Exp-Af: −0.98, Exp-Exp: −0.93, Af-Af: - 0.82. Right: Exp-Af: −0.96, Exp-Exp: −0.45, Af-Af: 0.82.

## Conclusions

Homology models can now be produced quickly for nearly any size or sequence of protein. The accuracy of backbones for AlphaFold structures can be close to that obtained from experimental structure which suggests they could be used in place of experimental structures for tasks such as virtual screening of drugs. It is therefore important to establish a threshold that informs when these docking studies would be expected to produce the same results on a homology model as if they were performed on an experimental structure. Establishing this threshold is especially challenging because docking can produce different results from an identical starting point of molecular and protein atomic coordinates. Here we avoid the uncertainty that could result from looking at a single docking comparison by evaluating many molecules with a wide range of sizes and properties.

Docking is typically used to explain in greater detail information that is already known from experimental studies. But the tool of docking can be used more generally as a means of characterizing the shape and interfacial properties of a protein without any intent to connect those results to experimental information. While docking is imperfect, we show here that deviations in its degree of reproducibility can be revealing in itself.

Starting with a large and highly varied set of molecules had the benefit of exploring many possible binding interactions. However, this large data set turned out to initially mask some of the most interesting observations. With the full set of molecules, no trend in median distance changes could be readily seen. However, when restricting the analysis to smaller sets of molecules with different numbers of rotatable bonds, the trends became more evident.

This study was possible because the CASP14 proteins all had experimental structures available. For AlphaFold models of novel proteins within a proteome there is no such available structure and no way to measure a GDT value. However, the pLDDT score have been shown to have a strong correlation to experimentally measured Cα-LDDT values. An examination of 10,215 recent experimental structures with <3.5 Å resolution revealed a Pearson correlation coefficient of 0.73 between pLDDT and Cα-LDDT.^11^ pLDDT appears to be a good surrogate for both Cα-LDDT and for GDT and we anticipate that the trends we report for GDT would be similar if benchmarked against Cα-LDDT. An advantage of using CASP structures in this study is that the experimental structures are not released until after the competition so the AlphaFold calculation could not be influenced by knowledge of those structures, whereas analysis of current AlphaFold structures could suffer from that issue.

The fact that more flexible molecules were particularly well-suited for identifying differences suggests a method for optimizing homology structures. Assessing difference in binding of flexible molecules during training could be used to classify those steps that improve the quality of a model as a docking template.

## Data Availability

All the software used, and all data sets are open source and freely available. The code used to run the docking, measure the distances, and create the figures is all available at: https://github.com/alplon/APHMDR_AP-SR

