## Supporting Info for "Assessing Protein Homology Models with Docking Reproducibility"


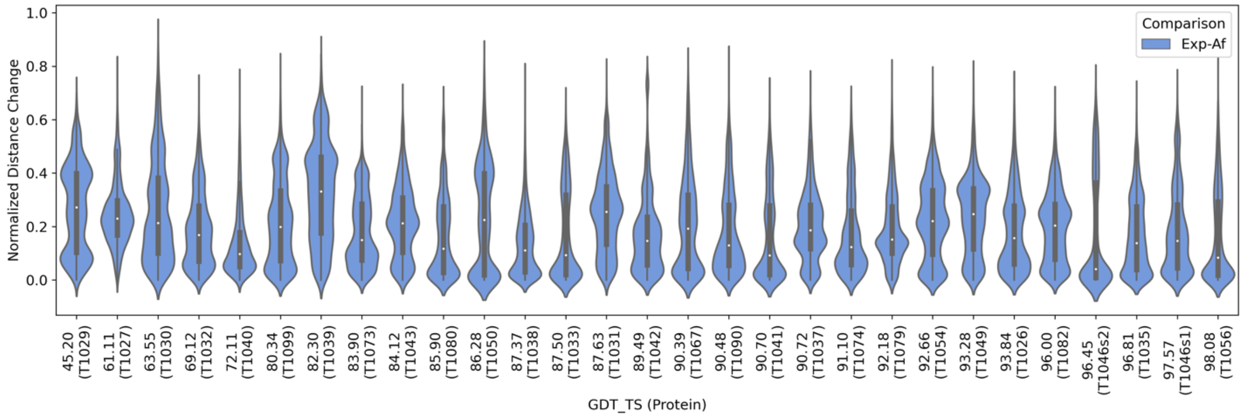


**Figure S1.** Normalized cross-comparisons of docking distance changes between docking to AlphaFold structures compared to docking to experimental structures displayed as violin plots. (Blue).

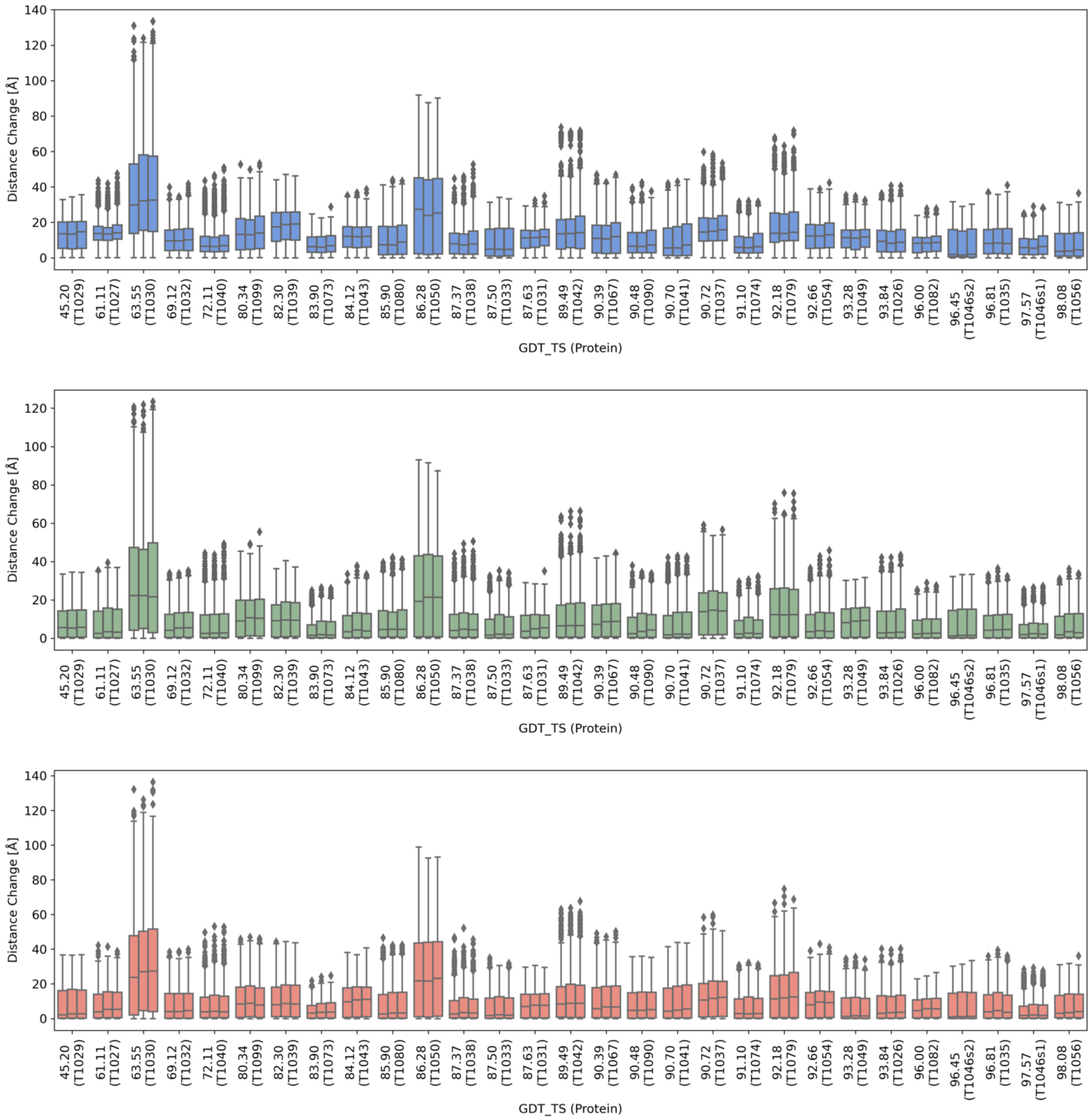


**Figure S2.** Non-normalized distance changes for cross- (top), and self-comparisons: Exp-Exp (middle) and Af-Af (bottom) separated by trial comparisons.

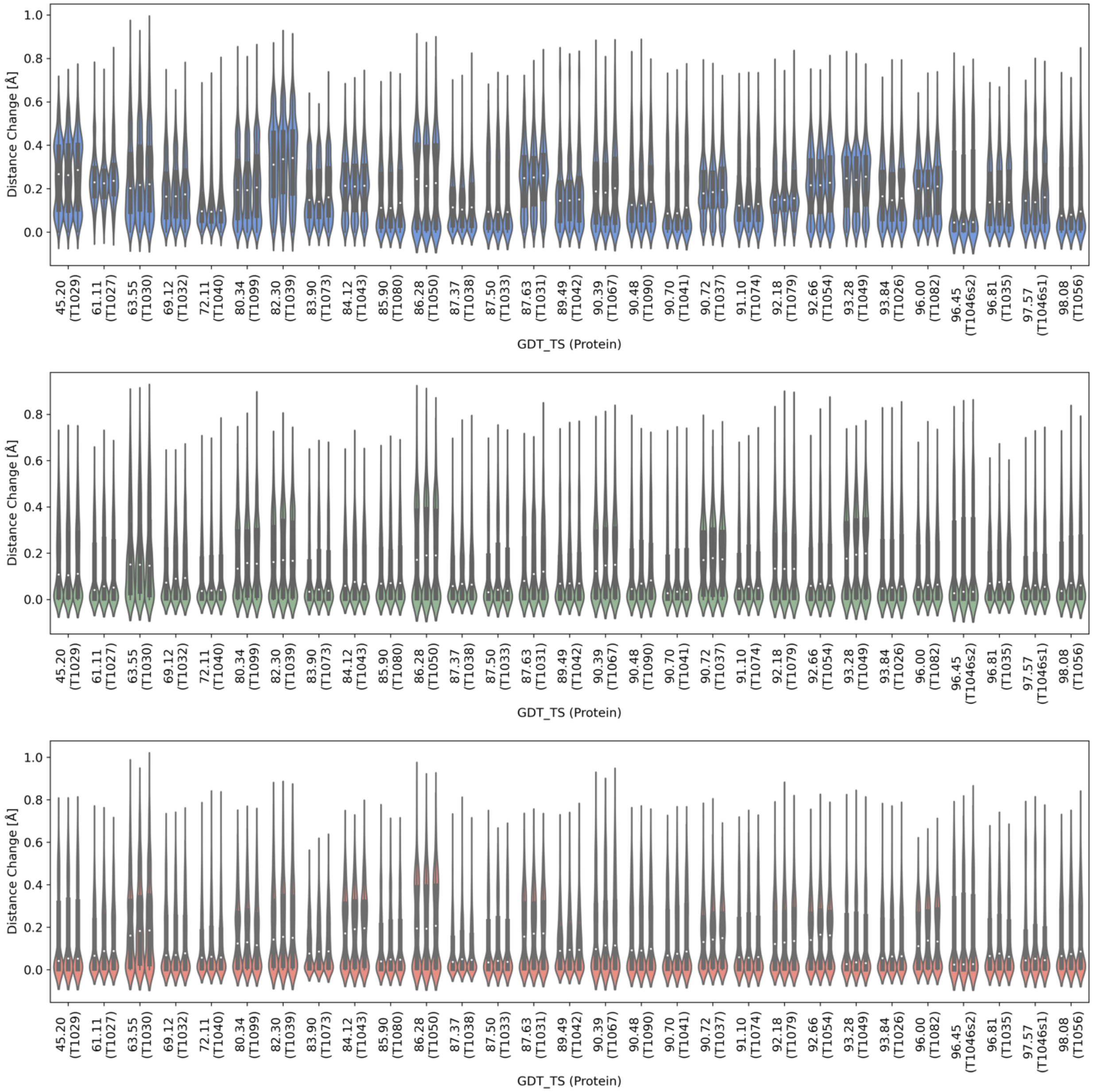


**Figure S3.** Normalized distance changes for cross- (top), and self-comparisons: Exp-Exp (middle) and Af-Af (bottom) separated by trial comparisons displayed as violin plots.

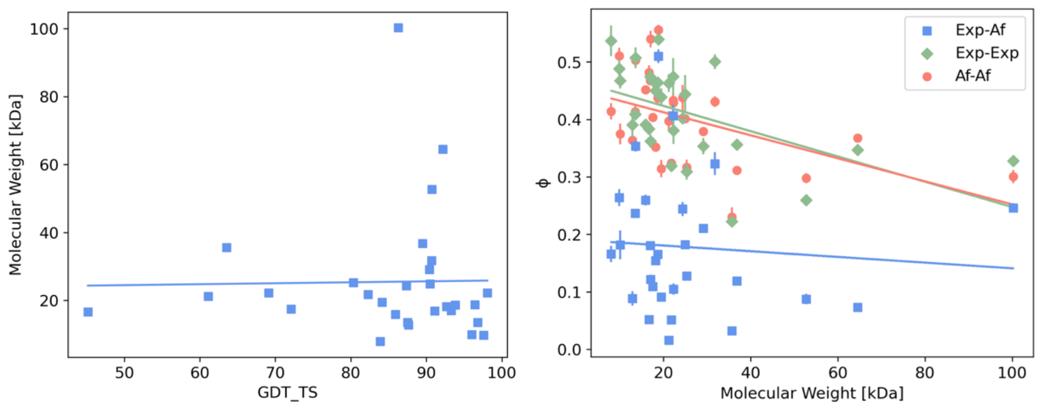


**Figure S4.** Left**:** Scatter plot of protein molecular weight and GDT_TS with a Pearson correlation coefficient of 0.02. Right: Scatter plot of protein molecular weight and fraction of molecules within 2 Å. The Pearson correlation coefficients are: Exp-Af: -0.10, Exp-Exp: -0.49, Af-Af: -0.49 (Red = AlphaFold-AlphaFold self-comparison, Green = experimental-experimental self-comparison, Blue = AlphaFold-experimental cross-comparison).

**Tables S1.** GDT_TS and molecular weights for each target.

| target | GDT_TS | molecular weight [kDa] |
|---|---|---|
| T1029 | 45.2 | 16.65 |
| T1027 | 61.11 | 21.22 |
| T1030 | 63.55 | 35.63 |
| T1032 | 69.12 | 22.28 |
| T1040 | 72.11 | 17.50 |
| T1099 | 80.34 | 25.28 |
| T1039 | 82.3 | 21.79 |
| T1073 | 83.9 | 7.96 |
| T1043 | 84.12 | 19.47 |
| T1080 | 85.9 | 15.89 |
| T1050 | 86.28 | 100.25 |
| T1038 | 87.37 | 24.34 |
| T1033 | 87.5 | 13.55 |
| T1031 | 87.63 | 12.85 |
| T1042 | 89.49 | 36.80 |
| T1067 | 90.39 | 29.10 |
| T1090 | 90.48 | 24.94 |
| T1041 | 90.7 | 31.78 |
| T1037 | 90.72 | 52.71 |
| T1074 | 91.1 | 16.92 |
| T1079 | 92.18 | 64.50 |
| T1054 | 92.66 | 18.19 |
| T1049 | 93.28 | 17.00 |
| T1026 | 93.84 | 18.65 |
| T1082 | 96 | 10.01 |
| T1046s2 | 96.45 | 18.78 |
| T1035 | 96.81 | 13.50 |
| T1046s1 | 97.57 | 9.84 |
| T1056 | 98.08 | 22.21 |

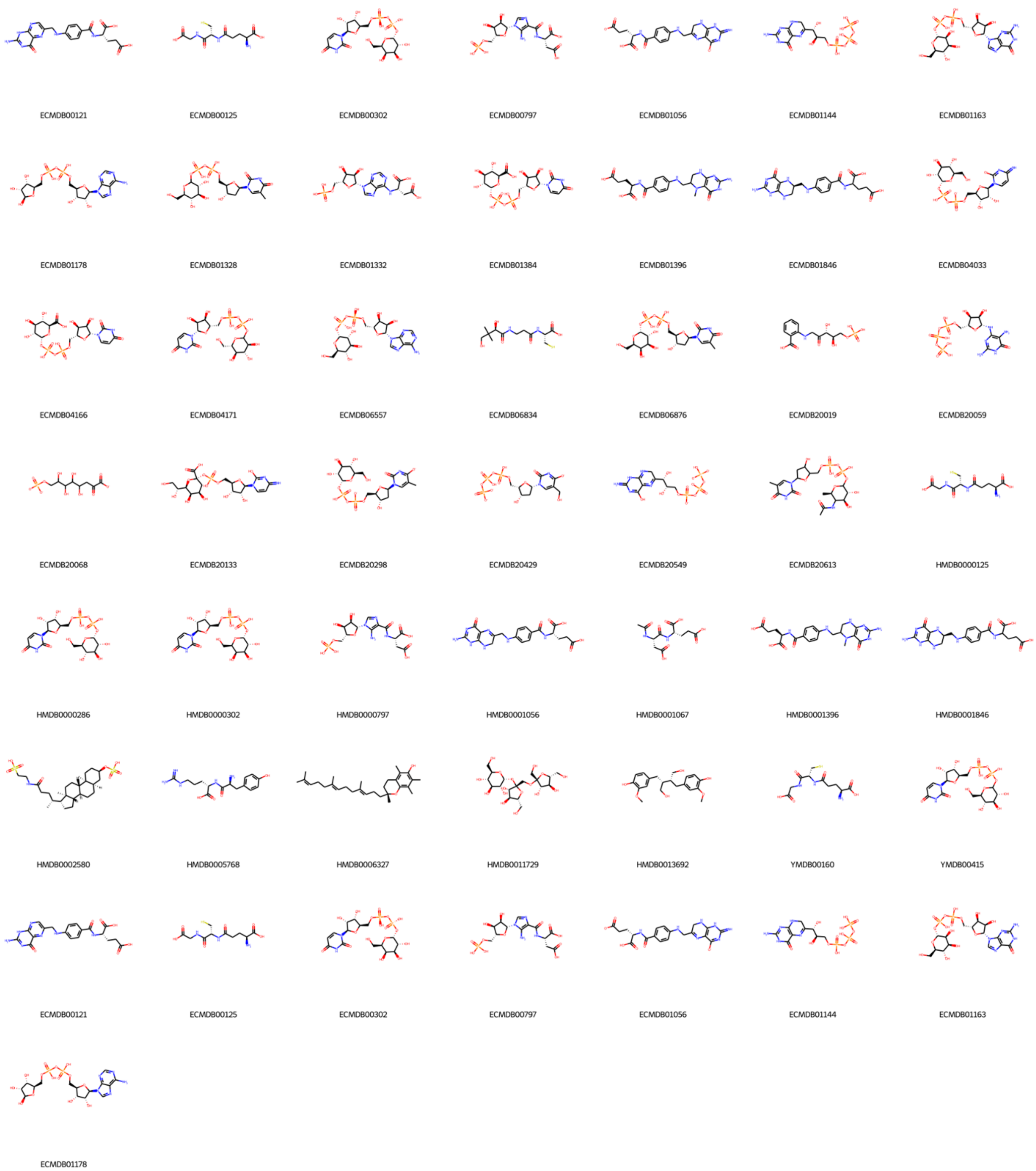


**Figure S5.** Molecular structures of all the probe subset with 9 rotatable bonds.

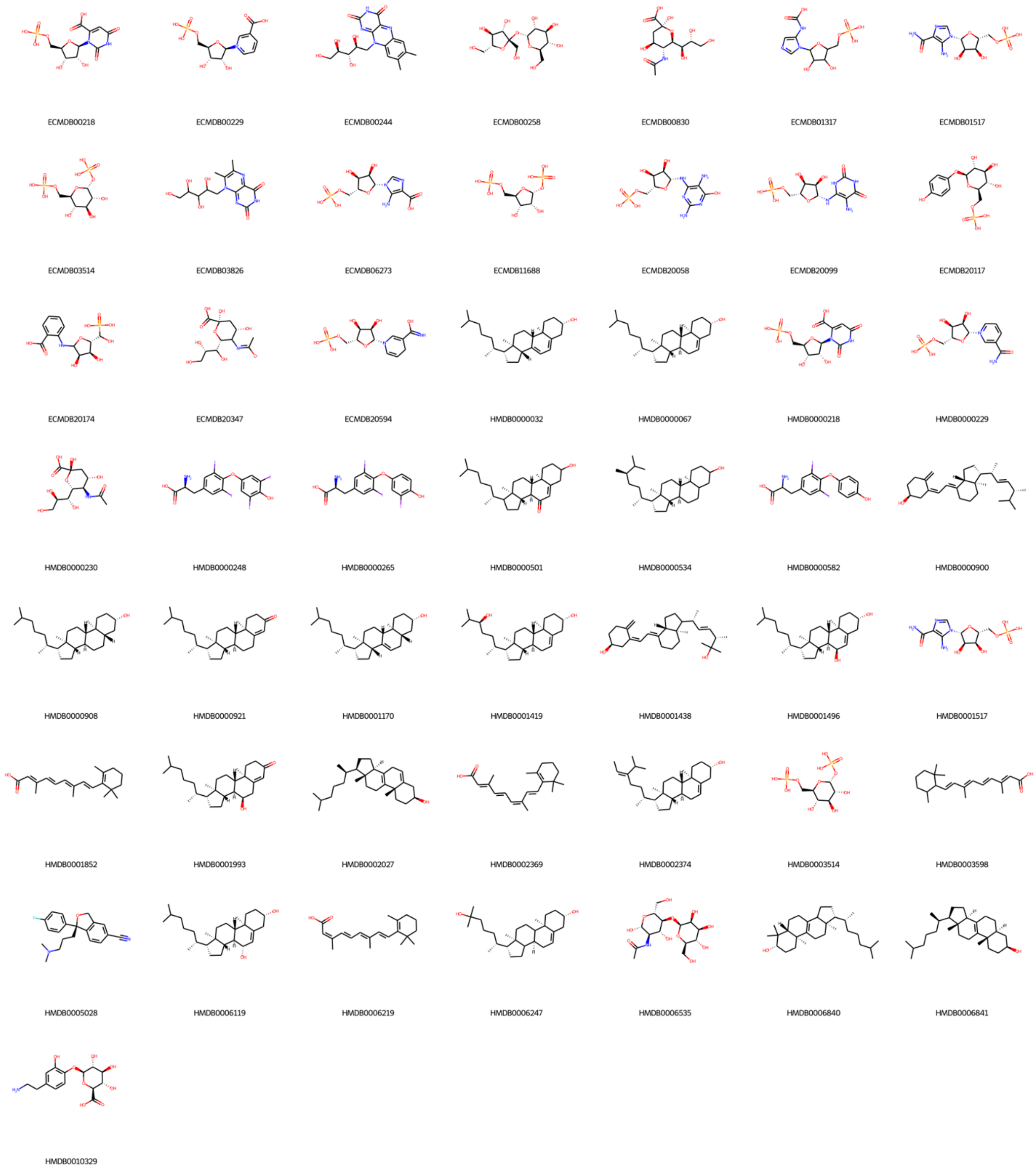


**Figure S6.** Molecular structures of all the probe subset with 5 rotatable bonds.

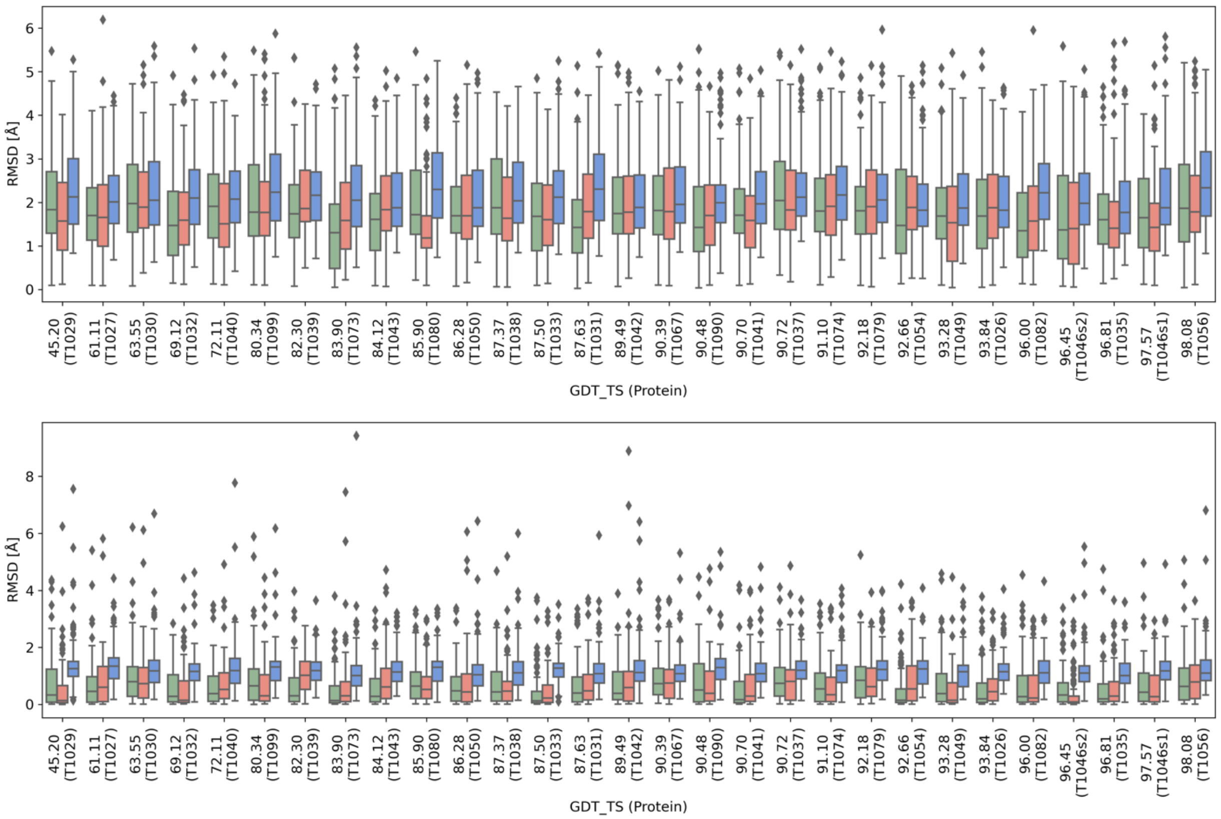


**Figure S7.** Boxplot of changes in RMSD for the placement of the molecules for self-comparisons and cross-comparisons for the probes with 9 (top) or 5 rotatable bonds (bottom) (Red = AlphaFold-AlphaFold self-comparison, Green = experimental-experimental self-comparison, Blue = AlphaFold -experimental cross-comparison).
